# Bacterial glycogen provides short-term benefits in changing environments

**DOI:** 10.1101/841718

**Authors:** Karthik Sekar, Stephanie M. Linker, Jen Nguyen, Alix Grünhagen, Roman Stocker, Uwe Sauer

## Abstract

Changing nutritional conditions challenge microbes and shape their evolutionary optimization. Here we investigated the role of glycogen in dynamic physiological adaptation of *Escherichia coli* to fluctuating nutrients following carbon starvation using real-time metabolomics. We found significant metabolic activity remaining after the depletion of environmental glucose that was linked to a rapid utilization of intracellular glycogen. Glycogen was depleted by 80% within minutes of glucose starvation and similarly replenished within minutes of glucose availability. These fast timescales of glycogen utilization correspond to the short-term benefits that glycogen provided to cells undergoing various physiological transitions. Cells capable of utilizing glycogen exhibited shorter lag times than glycogen mutants when starved between different carbon sources. The ability to utilize glycogen was also important for the transition between planktonic and biofilm lifestyles and enabled increased glucose uptake during pulses of limited glucose availability. While wild-type and mutant strains exhibited comparable growth rates in steady environments, mutants deficient in glycogen utilization grew more poorly in environments that fluctuated on minute-scales between carbon availability and starvation. Altogether, these results highlight an underappreciated role of glycogen to rapidly provide carbon and energy in changing environments, thereby increasing survival and competition capabilities in fluctuating and nutrient poor conditions.

## Introduction

Microbes must adapt to and compete under changing nutrient conditions. Instead of a well-mixed environment, bacteria in the wild often experience a feast-or-famine existence. Many microbial habitats are characterized by longer periods of nutrient starvation, intermittently punctuated by nutrient availability (Stocker, 2012). Thus, microorganisms face strong selective pressure to quickly resume growth when nutrients once again become available, and a diversity of strategies has evolved (Bergkessel, Basta, & Newman, 2016; Shoemaker & Lennon, 2018). Generally, these strategies involve the accumulation of unused resource that are labile and quickly activated when richer nutrient environments permit fast growth. For example, *Escherichia coli* facilitate rapid physiological transitions to higher quality nutrient conditions by maintaining a pool of ribosomes that only become translationally active as available nutrient becomes more abundant (Kohanim et al., 2018; Li et al., 2018; Metzl-Raz et al., 2017; Mori, Schink, Erickson, Gerland, & Hwa, 2017). *E. coli* also often feature additional enzymatic capacity beyond that immediately required (Davidi & Milo, 2017; O’Brien, Utrilla, & Palsson, 2016; Sander et al., 2019), and accumulate metabolically costly amino acids from protein degradation during starvation, which are then rapidly used for RNA and protein synthesis upon the resumption of growth (Link, Fuhrer, Gerosa, Zamboni, & Sauer, 2015). Strategies in other organisms include accumulation of alanine dehydrogenase in *Bacillus subtilis* to expedite growth after shifts to different environments (Mutlu et al., 2018), and the accumulation of methane oxidases in the methanotroph, *Methyloprofundus sedimenti*, induced by starvation in an effort to rapidly convert the next available methane into methanol (Tavormina et al., 2017).

Glycogen, a polymer of glucose, is another stored resource across evolutionarily divergent species. While the role of glycogen in mammalian cells is well-established as a temporary sugar reserve, the role of glycogen in bacteria such as *E. coli* has been less clear. Earlier studies have linked glycogen with long-term survival, contributing an energy source when the environment does not (Wilson et al., 2010); whereas, others discuss it as a temporary resource used during the physiological transitions necessitated by dynamic environmental conditions (Morin et al., 2017; Seok et al., 1997; Yamamotoya et al., 2012). Some studies combine the two perspectives, describing a role for glycogen that contributes to survival or maintenance in environments that frequently fluctuate in nutrient availability (Bourassa & Camilli, 2009; Jones et al., 2008). The concept of glycogen as a nutrient “bank” from which cells withdraw and deposit (Bertrand, 2019) summarizes the prevailing view about the role of glycogen in bacteria; however, for how long after starvation glycogen continues to supply the cell and towards what physiological processes it is used remains to be clarified.

Here, we describe the temporal dynamics of glycogen synthesis and breakdown between periods of nutrient availability and starvation. Using real-time metabolomics (Link et al., 2015) and glycogen measurements, we discovered that glycogen is depleted by more than 80% within 10 minutes of entry into starvation conditions and replenished after 2 min of nutrient availability. By comparing wild-type cells with cells unable to use glycogen, we found that glycogen shortens lag times when switching between carbon sources, enhances uptake when glucose is limited and facilitates the transition from planktonic to biofilm lifestyles. Importantly, this advantage conferred by glycogen existed only in dynamic or fluctuating environments; glycogen-deficient cells performed comparably to glycogen wild-type cells in steady environments. Our results suggest a role for glycogen during physiological transitions that involve starvation. We propose that glycogen serves as a short-term resource, consumed in the minutes after the onset of starvation. The short-term uses of glycogen may lead to long-term benefits; though from our data, it is unlikely that glycogen stores alone work to directly support bacterial maintenance in extended periods of nutrient starvation.

## Results

### Cells utilize glycogen upon carbon starvation

To investigate the role of glycogen during starvation, we designed a real-time metabolomics experiment to compare the metabolic changes across a transition into starvation of *E. coli* wild-type and a mutant unable to utilize glycogen. Specifically, we harvested minimal medium mid-log phase cultures at an optical density of 600 nm (OD) of 0.8 by fast filtration (Rabinowitz & Kimball, 2007) and resuspended them in the same medium but with a limiting amount of glucose as the sole carbon source (Fig 1A). We designed the medium such that the culture would deplete all carbon within 30-40 min (Supplementary Information). Across the transition into starvation, we measured over 100 metabolites as the sum of extra- and intracellular molecules every 15 s using real-time metabolomics (Link et al., 2015). In wild-type cells, the ion corresponding to hexoses such as glucose was depleted within 30-40 min (Fig 1B). Several ions annotated to central carbon metabolites diminished immediately after glucose was depleted (Figure S1), but others such as hexose phosphate and amino acids remained stable or even increased such as the tricarboxylic acid (TCA) cycle intermediate (iso)citrate (Fig 1B). The large number of stable or even increased metabolites suggest ongoing metabolism that is supplied from another source. Given the stable concentration of hexose phosphates and that the first step of glycogen hydrolysis releases glucose-1-phosphate, we hypothesized that glycogen usage may supply metabolism. Indeed, by performing the same experiment with the *glgP* mutant that is unable to use glycogen, we observed a similar depletion of glucose across the shift into starvation. In contrast to the wild-type, however, the level of hexose phosphates in the *glgP* mutant depleted concurrently with glucose (Fig 1B). Additionally, other metabolite levels were reduced compared to the wild-type (Fig 1B). Glycogen utilization did not explain stable levels of all ions during transition to starvation (Figure S1B); specifically, the abundances of ions corresponding to metabolites 3-propylmalate, isopropylmaleate, and orotate remained roughly constant in both strains. Nonetheless, the depletion of hexose phosphates in the wild-type versus *glgP* strain implicates the utilization of glycogen within minutes of the transition into starvation.

**Figure 1.**
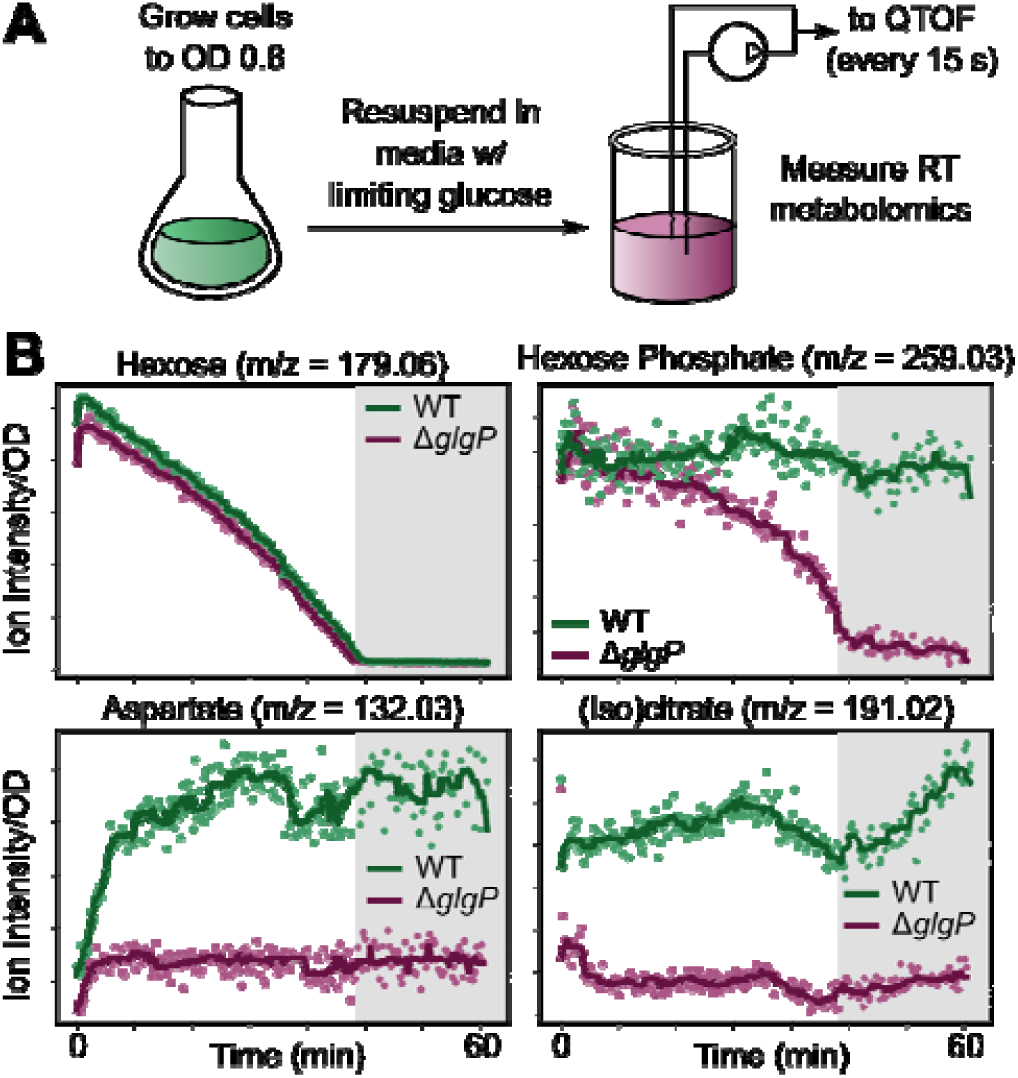
Cells show secondary, glycogen-related metabolic activity upon carbon starvation. (**A**) Experimental setup for measuring metabolic profile of cells depleting carbon. Growing cells were switched to medium with limiting glucose (0.32 g/L), then real-time metabolomics (Link et al., 2015) was measured for a total of 1 h. For real-time metabolomics measurement, a pump circulated culture and injected 2 μL of culture directly into a quantitative time of flight mass spectrometer every 15 s. (**B**) Glycogen mutant cells observe different metabolic activity on transition to starvation. Traces of exemplary ions are shown that correspond to hexose, hexose phosphate, aspartate, and (iso)citrate for two strains, WT (wild-type, green) and a *glgP* mutant (purple). Dots indicate ion intensity measurement normalized to initial OD. Gray area indicates the time period after glucose depletion. Solid lines are a moving average filter of the measured ion intensity.

To test the hypothesis that a rapid onset of glycogen breakdown serves as an immediate fuel, we measured cellular glycogen content from the onset of starvation to 50 min after starvation (Figure 2A). We found that glycogen content diminished by 80% within the first 10 min of starvation. Thus, *E. coli* consumes glycogen rapidly after carbon depletion, potentially enabling the pronounced metabolic activity we observed even hours after starvation entry (Figure 1). To elucidate how rapidly the glycogen storage is replenished upon the return of carbon availability, we added fructose to a culture that was carbon starved for 30 min. Fructose was chosen because glucose supplementation would have interfered with the ability to accurately measure glycogen content. Upon fructose addition, the intracellular glycogen content reached a steady glycogen level within 2 min (Figure 2B). Thus, glycogen synthesis and degradation occur on minute time scales, suggesting that glycogen serves a potential role as a short-term energy storage in microbes, akin to the mammalian system.

**Figure 2.**
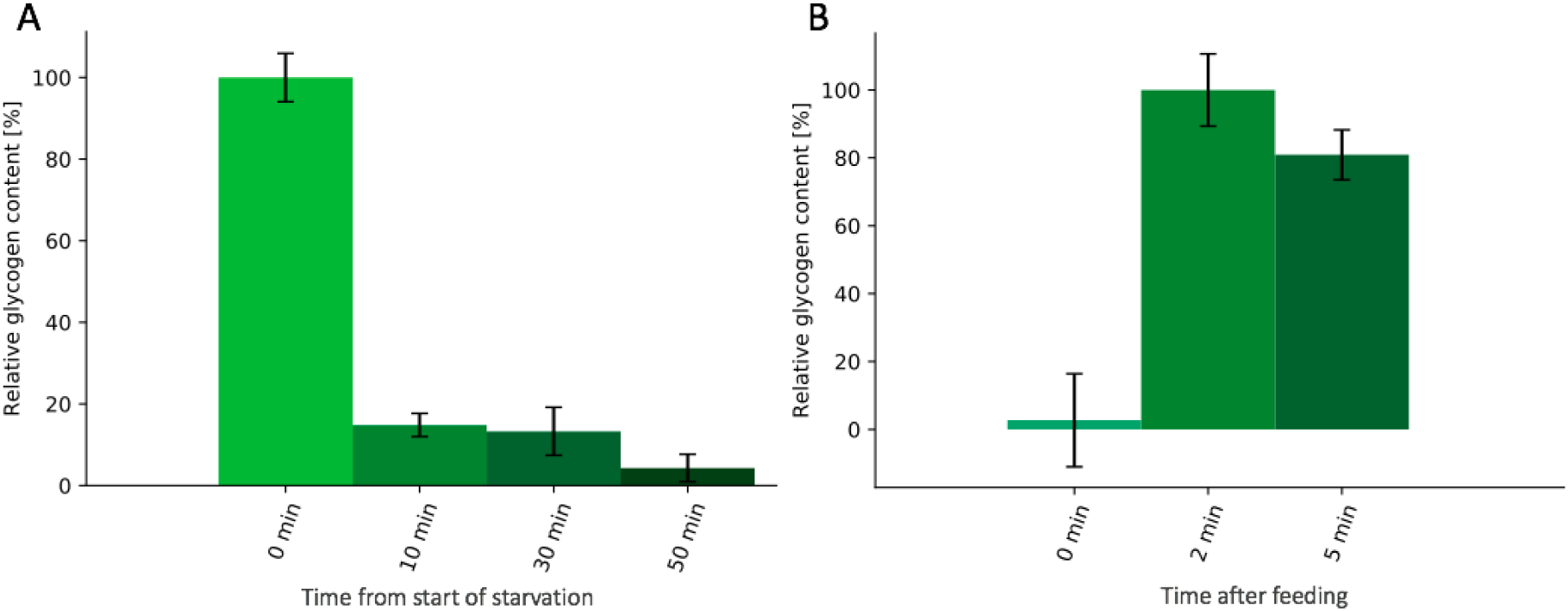
Intracellular glycogen depletes rapidly after carbon downshift and accumulates rapidly on carbon availability. **(A)** Glycogen depletion during starvation. Wild-type cells were grown to mid-log phase (OD 0.8) and resuspended in medium without carbon. The first sample time point was taken before resuspension. Error bars indicate the standard error of 6 biological replicates. **(B)** Rapid glycogen synthesis upon fructose addition to a carbon starving culture. Fructose was added at time point zero to wild-type *E. coli* harvest from mid-log phase (OD 0.8) after 30 min starvation in medium without carbon. Error bars indicate the standard error of 3 biological replicates.

To elucidate the complete dynamics of the metabolic response to glycogen degradation and synthesis, we designed a more controlled real-time metabolomics experiment. Specifically, we fed glucose to a culture that was starved for 30 min at a constant rate of 8 mmol glucose/g dry weight/h for 5 min, then we turned off the feed pump, and we measured metabolism for an additional 80 min. The feedrate of 8 mmol/g/h was chosen to be well below the maximum uptake rate of *E. coli* (Monk et al., 2016; Sekar et al., 2018), meaning that glucose will not abundantly accumulate in the medium. Consistent with this design, the ion corresponding to glucose depleted within 1-2 min after the feed ceased (Figure S2). We observed a sudden drop after glucose depletion in all other metabolite concentrations including hexose-6-phosphate, (iso)citrate, and other central carbon metabolites for both wild-type and *glgP* (Figure 3). In contrast to the *glpP* mutant, several metabolites within or near the TCA cycle exhibited a secondary response in the wild-type. After initial depletion, isocitrate, in particular, immediately arises again within 5 min to a level near that of the glucose fed state. This “bounce” effect was also observed prominently in glutamine, glutamate, malate, and aspartate, as indicated by the green arrows. That the bounce effect was observed primarily in metabolites within or near the TCA cycle (Figure 3, Figure S3) suggests that glycogen is used to fuel respiration right after the onset of starvation.

**Figure 3.**
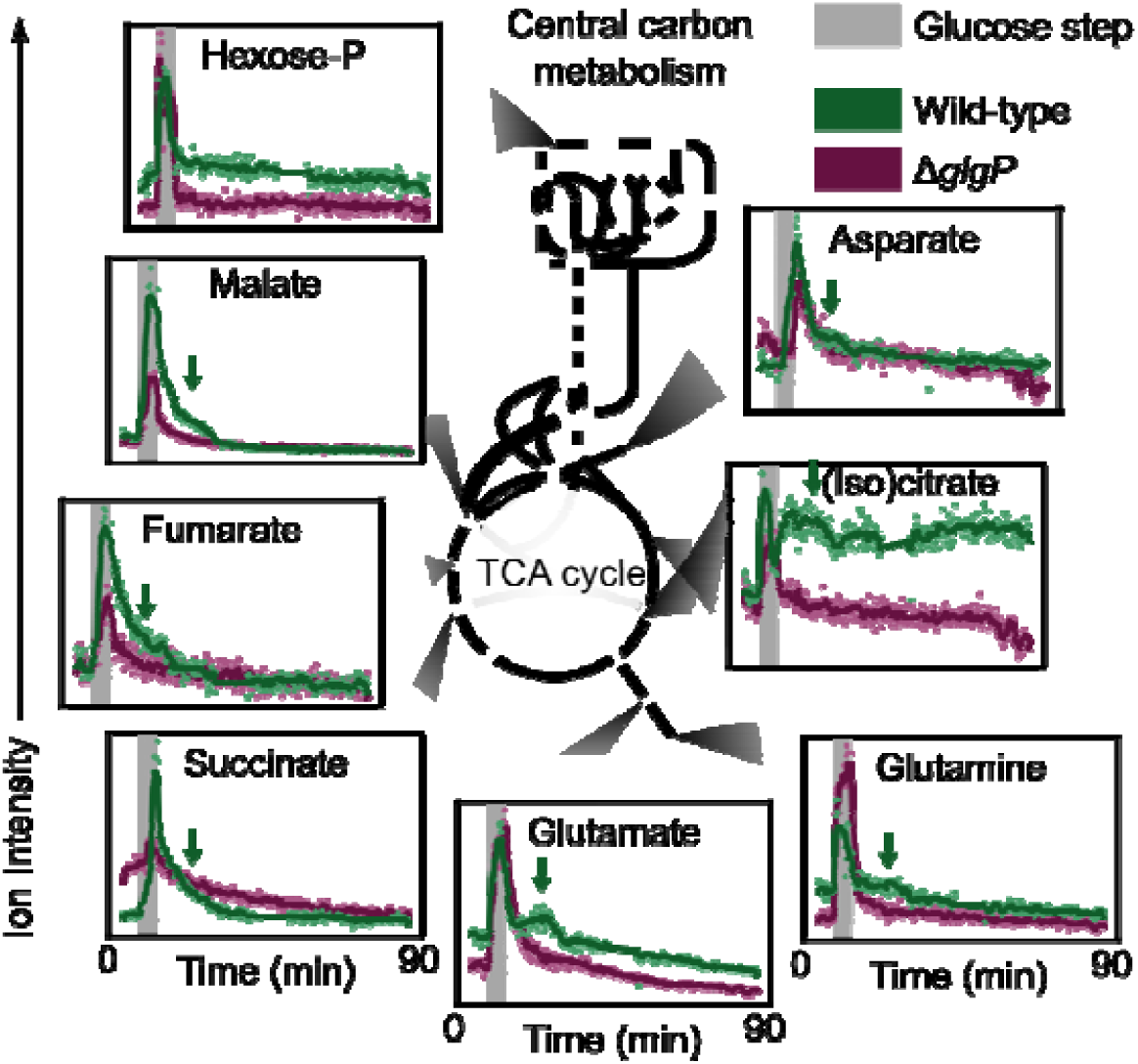
A glycogen-related metabolic response occurs in response to brief, constant glucose feed. *E. coli* was grown to OD 0.8-1.2, starved for 30 min without glucose and fed a constant glucose supply. The glucose was supplied with a pump at a rate of 8 mmol/g/h for two strains, WT (wild-type, green) and a *glgP* mutant (purple). After 5 min of glucose application, the glucose feed was ceased. Throughout the feed, real-time metabolomics measurement was performed, and data is shown for ions corresponding to central carbon metabolites. The green arrows indicate a metabolic “bounce” where measured ion intensity increases 5-15 minutes for metabolites malate, fumarate, succinate, glutamate, glutamine, (iso)citrate, and asparatate. Dots indicate ion intensity measurement normalized to initial OD. Gray area indicates the time period where glucose was supplied. Solid lines are a moving average filter of the measured ion intensity.

Overall, we posit that glucose starvation initiates glycogen utilization, both during gradual glucose depletion as in the earlier experiment or the nearly instantaneous depletion here. These observations are consistent with known and suggested interactions of glycogen phosphorylase and glucose uptake-related proteins (Seok et al., 1997; Tian, Fauré, Mori, & Matsuno, 2013); specifically, the HPr protein involved in glucose uptake positively activates glycogen phosphorylation allosterically. A strongly stimulatory effect occurs when HPr is dephosphorylated as is typical for starvation. The rapid timescale enabled by allosteric regulation is consistent with our data, which suggests that decreasing glucose uptake rapidly triggers glycogen usage.

### Glycogen grants advantage in changing conditions

The minute-scale liquidation of glycogen led us to ask whether glycogen enables cells to accommodate sudden environmental change. To evaluate how glycogen affects the ability to adapt to new environments, we tested two biologically relevant transitions: a change of nutrient source and the transition from planktonic to biofilm growth. As a control, we first tested the influence of glycogen in stable environments and determined that the difference in the steady-state growth rate of wild-type cells versus different glycogen mutants was small (within 15%; Figure 4A). Next, we performed a nutrient-shift experiment where wild-type and *glgP* mutant were grown to mid-log phase (OD 0.4) in glucose medium. After centrifugation and washing, cultures were rapidly transferred into a medium with acetate as the sole carbon source, either directly or with a transitory 30 min period of starvation in carbon-free medium. Without starvation, the time to resume full growth after the switch (i.e. the lag time) was identical for wild-type and mutant (Figure 4B). With an intermittent starvation period, however, the glycogen mutant exhibited a roughly doubled lag time (∼220 min versus ∼110 min) with respect to the wild-type. To test whether this reliance on glycogen was also required during less abrupt transitions, we performed a modified lag time experiment, where acetate was added either 60 min before or 60 min after glucose was depleted from the initial medium (Figure 4C). Consistent with the previous experiment, we found comparable lag times between the glycogen mutant and wild type without starvation. However, after a period of starvation, the lag time of the glycogen mutant was again significantly prolonged with respect to the wild-type. Presumably, the wild-type has a shorter lag time after starvation because they either initiate the adaptation already before depletion of the primary carbon source or scavenge previously excreted carbon sources such as acetate (Mandel & Silhavy, 2005; Rahman, Hasan, Oba, & Shimizu, 2006; Wei, Shin, LaPorte, Wolfe, & Romeo, 2000). Our data suggests that cells unable to use glycogen are consequently slower in completing the necessary molecular adaptions for full growth in new conditions. Likely, these cells are deprived of alternative carbon and/or energy sources when experiencing a change in carbon source.

**Figure 4.**
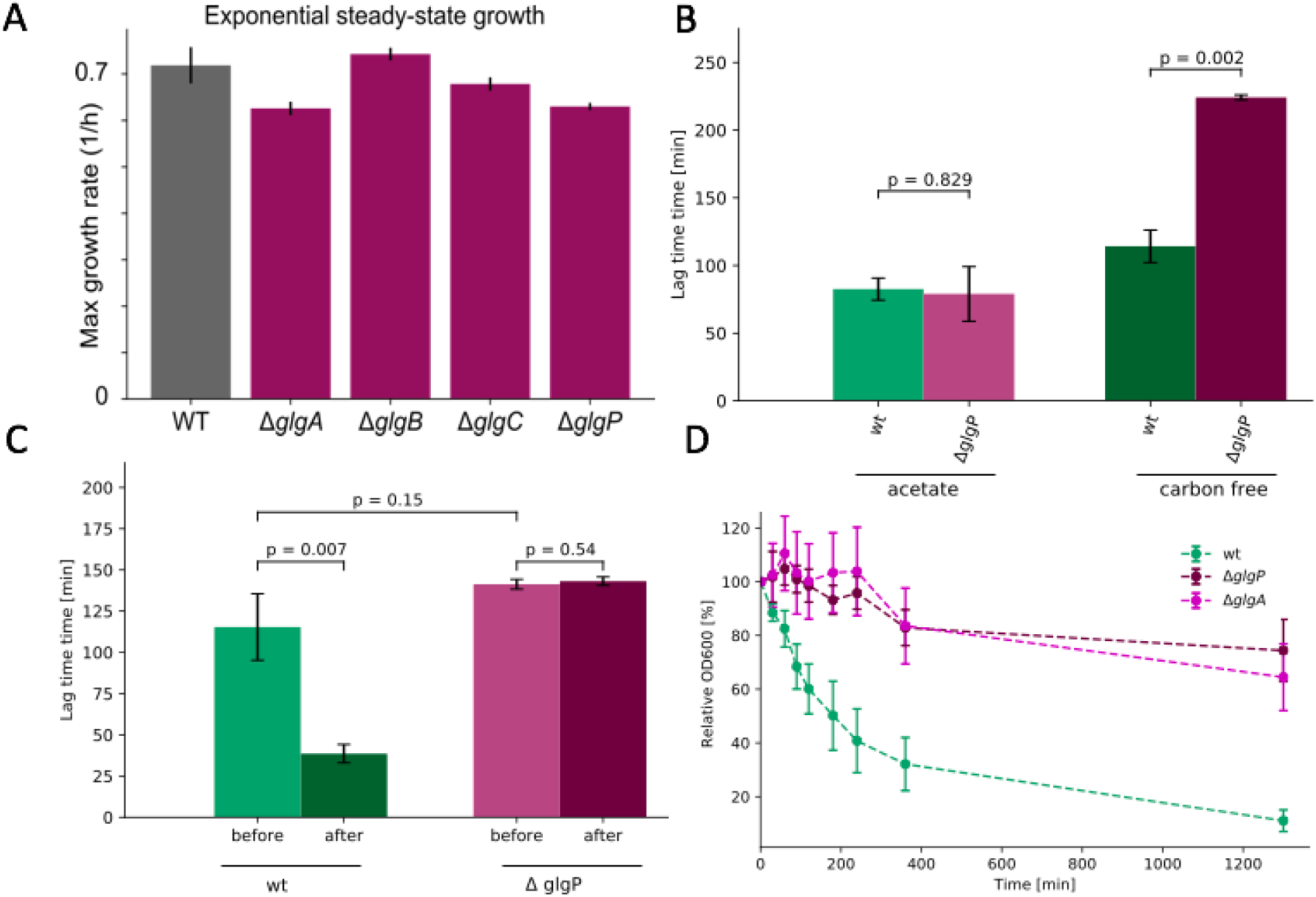
Glycogen-related phenotypes in steady state versus changing conditions. (**A**) Glycogen mutants exhibit similar growth rates to wild-type under steady-state growth. (**B**) The *glgP* glycogen mutant exhibits prolonged lag times when starved between nutrient transitions. Wild-type and *glgP* mutant cells were grown to mid-log phase (OD 0.4) in glucose media. Cells were rapidly transferred into acetate medium either directly or with in-between 30 min period of starvation in carbon-free medium. The lag time until growth resumption was measured for all cells. Error bars indicate the standard error of 3 biological replicates, and P values were calculated assuming independence with Student’s t test. **(C)** Wild-type and *glgP* mutant cells were grown to mid-log phase (OD 0.4) in glucose media. Cells were rapidly transferred into acetate medium either 60 min before or 60 min after glucose depletion in the initial media. The lag time until growth resumption was measured for all cells. (**D**) Glycogen mutants remained planktonic in stationary phase. Wild-type, *glgP*, and *glgA* mutant cells were grown until stationary phase. Afterwards, cells were cultivated without shaking to initiate biofilm formation. Cell attachment was measured via optical density.

The transition from planktonic to sessile (biofilm) lifestyles represent another adaptation that requires substantial restructuring of cellular physiology. Biofilm formation is characterized by three phases: attachment, maturation, and dispersal (Weiss, Obied, Kalkman, Lammertink, & van Leeuwen, 2016). We focused on the attachment phase, which is characterized by the decrease of planktonic cells. A common method for estimating the concentration of planktonic bacteria relies on measuring the OD_600_. When stationary phase *E. coli* was cultured without shaking, the number of planktonic cells decreased by 89% within 18 hours (Figure 4D). The *glgP* and *glgA* mutants, in contrast, remained largely planktonic even after 18 hours (26% and 36% decrease, respectively). Therefore, wild-type cells have either an increased attachment rate or an increased mortality rate in comparison to the glycogen mutant. The latter is unlikely as our previous experiments have indicated metabolic, viable activity for cells well into starvation. Biofilm formation is induced by nutrient starvation and inhibited by glucose addition (Thomason, Fontaine, De Lay, & Storz, 2012; Zhao et al., 2017). We therefore reason that glycogen facilitates the attachment phase of biofilm formation under starvation conditions, here by providing resources for matrix protein or flagella production.

### Glycogen utilization confers a growth advantage in dynamic nutrient environments

Given the importance of glycogen during physiological transitions, we sought to establish the growth advantage conferred by glycogen utilization under controlled, dynamically changing conditions. By coupling microfluidics and time-lapse imaging, we monitored the volumetric growth of individual *E. coli* cells under fluctuating and steady nutrient supply. The fluctuating environment consisted of 30 s long nutrient pulses followed by 5 min of carbon starvation, whereas in the steady environment the carbon source was continuously replenished (Figure 5A). In both environments, precise control over the nutrient signal was maintained by flowing medium over surface-attached cells and switching between two media when generating a pulse (Nguyen, Fernandez, et al., 2019; Sekar et al., 2018). In these environments, we competed the YFP-labeled wild-type and the CFP-labeled *glgP* mutant and monitored their growth through image analysis.

**Figure 5.**
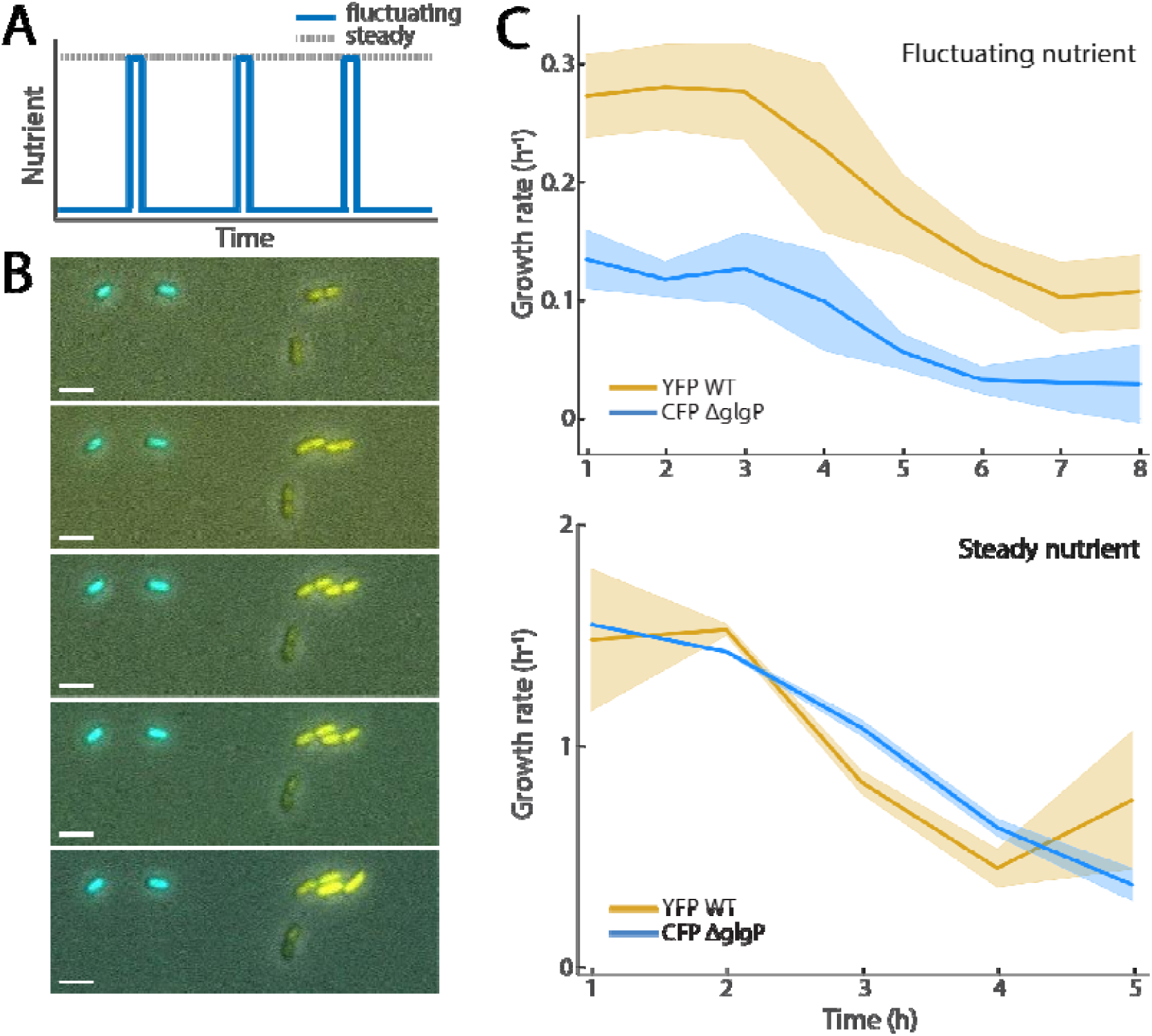
Glycogen consumption offers a growth advantage in pulsing nutrient environments. (**A**) Nutrient signals in pulsing and steady environments for microfluidic experiments. Low phases of the fluctuating signal (blue) deliver zero carbon for 5 min, while the high phases are 30 second pulses of 2% (v/v) LB solution. The steady nutrient maintains a constant concentration of 2% LB solution. (**B**) Montage of composite images from three channels: phase (grayscale), YFP (yellow) and CFP (cyan). Scale bar indicates 3 μm. (**C**) Growth rate over time from wild-type and Δ*glgP* populations within pulsing and steady environments. In both panels, curves represent the mean growth rate across replicate experiments; error bars represent standard deviation between replicates (three biological replicates across separate days for pulsing condition; two biological replicates across separate days for steady). Each replicate observed at least 1167 individual *E. coli*.

In fluctuating environments, cells capable of consuming glycogen had an apparent growth advantage over those that could not. From time-lapse images, YFP-labeled wild-type cells visibly increased in cell mass and often divided, while the CFP-labeled *glgP* mutant hardly grew in size (Figure 5B). We then quantified single-cell growth rate as the rate at which cell volume exponentially doubles, as assessed from image frames captured 3 min apart. In fluctuating environments, these quantifications yielded maximum specific growth rates of 0.28 ± 0.04 h^−1^ and 0.13 ± 0.03 h^−1^ for wild-type and mutant strains, respectively, whereas in steady environments, the maximum specific growth rates of the two strains were indistinguishable (Figure 5C). To summarize, the ability to utilize glycogen enhances growth in fluctuating environments, thereby substantiating a key role for glycogen as an immediately available resource across changing environments.

### Glycogen utilization enable improved nutrient uptake capability

So far, we have established that glycogen utilization confers a growth advantage in dynamic environments by providing energy and carbon in nutrient poor transition phases. It is not clear which cellular functions are supplied by the freed carbon from liquidated glycogen beyond biofilm faculties. Nevertheless, we hypothesized that the at least some of the freed carbon would lead to better uptake ability, a paramount survival attribute in scant environments. To measure the cellular ability for nutrient uptake, we used real-time metabolomics to monitor glucose uptake while switching the cells between starvation and pulses of glucose (Sekar et al., 2018). As in the antecedent study, we observed rapid assimilation of glucose, as indicated by the detected levels of the ion corresponding to glucose (Figure 6A). Each pulse showed an instantaneous increase of glucose concentration followed by depletion caused by bacterial consumption. Fitting a Michaelis-Menten model to the glucose consumption, where the uptake rate equates to the *V*_max_ of the fit (Figure 6B), revealed a much lower maximum capacity for glucose uptake in the *glgP* mutant compared to the wild-type (Figure 6C).

**Figure 6.**
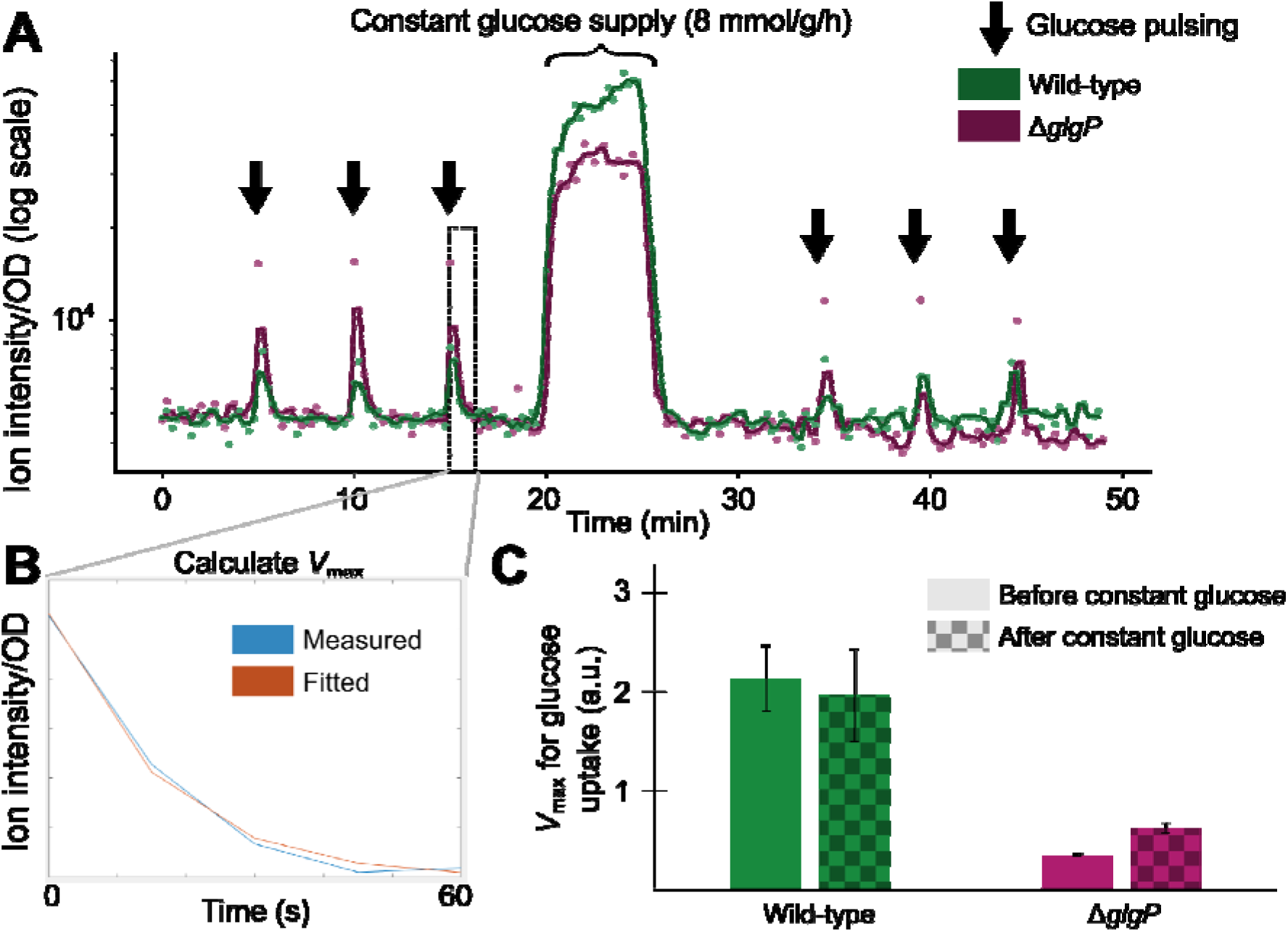
Glycogen capability enables increased scavenging in starvation. (**A**) Real-time metabolomics measurement of the ion corresponding to glucose in starved cells. Cells were grown to OD 0.8, then switched to media without carbon. Cellular metabolism was measured with real-time metabolomics as cells were pulse fed glucose every 5 minutes, an integrated feedrate of ∼0.4 mmol/g DCW/h (raw data available in **Supplementary Data**). After 20 min, a glucose was constantly supplied at 8 mmol/g/h for 5 minutes. After the constant glucose supply, cells were pulse fed glucose again every 5 min. (**B**) VThe kinetics of the glucose uptake were fitted to a Michaelis-Menten equation in order to calculate *V*_max_ for every pulse. (**C**) The calculated *V*_max_ (scavenging ability) is much lower for a *glgP* mutant compared to wild-type. The scavenging ability increases after the constant glucose supply for the *glgP* mutant; whereas, the WT strain’s scavenging ability is not improved. Error bars indicate the standard error of the glucose uptake rate for the pulses (*n* = 3). Dots indicate ion intensity measurement normalized to initial OD. Gray area indicates the time period after glucose depletion. Solid lines are a moving average filter of the measured ion intensity.

To test whether the difference in uptake capacity stemmed primarily from the carbon release in glycogen, we simulated the carbon release by providing a short dose of carbon by feeding glucose at 8 mmol/g/h for 5 min (Figure 6A), after the first set of limiting glucose pulses. Consistent with our hypothesis, the glucose uptake capacity remained high for the wild-type but improved significantly for the *glgP* mutant. Thus, the uptake capability does appear to originate from access to a nutrient source during starvation, whether its internal glycogen or additional carbon input. This carbon supply may fuel the synthesis of uptake related proteins, which are transcriptionally controlled by starvation related effectors (e.g. Crp) (You et al., 2013). The carbon supply may also prime the cells metabolically for carbon uptake, for example through high phosphoenolpyruvate (PEP) abundance. PEP is the substrate to phosphorylate incoming glucose through the phosphotransferase system, the primary means of rapid glucose uptake. While we did not measure PEP directly, we noticed differences in the energy charges, AMP and ADP, between wild-type and the glycogen mutant during starvation (Figure S4). Specifically, AMP and ADP were approximately 2.5 and 1.4 times more abundant in the mutant compared to wild-type, respectively. Difference in charge are often associated with changes in PEP abundance due to the dependence of PEP-associated carboxylases and kinases on the energy charges (Sauer & Eikmanns, 2005). In summary, glycogen release enables cells to perform more rapid uptake: an important capability when environments change often and nutrients are available only fleetingly.

## Discussion

From our findings, we propose a role for bacterial glycogen in dynamic environments. We found that glycogen is used to an appreciable magnitude in a short span of time (∼80% within 10 minutes), as glucose availability goes to zero. This demonstrates that glycogen is not merely a long-term energy storage that supplies microbial maintenance. Instead, glycogen is used within minutes for immediate physiological changes such as resumption of growth, induction of the attachment phase of biofilm formation, and to enable scavenging of nutrients. Furthermore, glycogen utilizing cells exhibited faster growth rates in dynamic environments, such as single nutrient shifts or repeated nutrient fluctuations, than glycogen-deficient cells. Altogether, our data reveals glycogen as a crucial internal resource, consumed within minutes of carbon starvation and synthesized within minutes of carbon re-availability, to aid in the physiological transitions that accompany environmental change.

Environmental change imposes physiological challenges to bacteria. For example, in nutrient-rich conditions, cells are not limited by their ability to scavenge nutrients. However, in starvation, the opposite is needed — cells must take up diverse nutrients much more efficiently (Towbin et al., 2017; You et al., 2013). The two scenarios result in a dilemma: the cell has a contrarian objective after switching between carbon rich and poor conditions. Meeting the new objective requires an appreciable change either in the abundance of key proteins for uptake or to reconfigure the cells metabolically (*e.g.*, elevated concentrations of PEP). Our data depict glycogen as a solution: a fast, flexible store of nutrients. While inability to use glycogen does not prevent cells from making physiological transitions, the ability to use glycogen seems to quicken the rate at which transitions occur. Thus, we show that glycogen is a facile resource for the cell to more quickly adjust its physiology to compete more effectively in starvation and nutrient poor conditions.

## Methods

### Strains and plasmids

*E. coli* BW 25113 from the Keio collection (Baba et al., 2006) was used as the wild-type (WT) strain for all experiments. Kanamycin markers were excised from the Keio knockout strains *glgP, glgA, glgB, and glgC* using pCP20 and verified using PCR (Datsenko & Wanner, 2000). All strains are listed in **Table S1** and plasmids are listed in **Table S2**. Strains and plasmids are available from authors on request.

### Cultivation, media, and real-time metabolomics profiling

Glucose media and culture preparation was followed as described in a previous study (Sekar et al., 2018). On the day before experiments, an inoculum of cells was prepared in sterile Luria-Bertani (LB) broth (10 g/L NaCl, 10 g/L bacto-tryptone, and 5 g/L yeast extract) in the morning and cultivated at 37°C with 225 RPM shaking until noon. At noon, cells were 1:50 diluted into M9 minimal medium + 0.4% glucose. In the evening, shake flasks with 35 mL of M9 medium + 0.4% glucose were prepared with 1:100 dilution from the M9 inoculum and cultivated at 30°C with 225 RPM shaking until the next morning. On the morning of the experiment, cells were typically OD 0.1 and then cultivated at 37°C with 225 RPM shaking until they reached OD 0.8, at which point the experiments were commenced. The M9 minimal medium consisted of the following components (per liter): 7.52 g Na_2_HPO_4_ · 2 H_2_O, 5 g KH_2_PO_4_, 1.5 g (NH_4_)_2_SO_4_, 0.5 g NaCl. The following components were sterilized separately and then added (per liter of final medium): 1 mL 0.1 M CaCl_2_, 1 mL 1 M MgSO_4_, 0.6 mL 0.1 M FeCl_3_, 2 mL 1.4 mM thiamine-HCL, and 10 mL trace salt solution. The trace salt solution contained (per liter) 180 mg ZnSO_4_ · 7 H_2_O, 120 mg CuCl_2_ · 2 H_2_O, 120 mg MnSO_4_ · H_2_O, 180 mg CoCl_2_ · 6 H_2_O. The real-time metabolomics profiling is fully described in (Link et al., 2015), but briefly: Cells were cultivated in a Schott bottle submerged in a water bath controlled at 37°C. Mixing and aeration were provided by a magnetic stirrer. A peristaltic pump circulated the culture through a six-port valve. On measurement, the valve configuration diverted roughly 2 μL of culture into a continuous flow of negative ionization buffer (60:40 vol/vol isopropanol:water with 1 mM ammonium fluoride, pH 9.0). The ionization buffer, now mixed with the live cells, was introduced for ionization in an electrospray chamber, and ions’ abundances were measured semi-quantitatively using a Quantitative Time of Flight mass spectrometry detector (Agilent 6550). Measurement (mixing of culture into the buffer) occurred every 15 seconds, thereby generating a time profile of the intracellular metabolic concentration. The annotation of ions is described in (Fuhrer, Heer, Begemann, & Zamboni, 2011).

### Real-metabolomics profiling of cells with depleting glucose

Cells were grown to mid-log phase where the optical density (OD) at 600 nm was measured to 0.8. At this point, 32.5 mL of the cells were collected on filter paper using fast filtration technique (Rabinowitz & Kimball, 2007) and rapidly resuspended into 25 mL of pre-warmed 1:8 diluted M9 medium (37 °C) with 0.32 g/L glucose as the sole carbon source within a Schott bottle. Immediately after resuspension, the real-time metabolomics profile of the cells was measured for 1 h.

### Lag phase experiments

To calculate the lag time of the glucose to acetate switch, cells were grown over night in M9 medium with glucose as carbon source at 37°C. The next day, cells were freshly inoculated in M9 medium with glucose and grown until OD 0.4. The cells were transferred in M9 medium with acetate either directly or with an intermediate starvation period of 30 min in carbon free media. For the transfer, the collected cells were rapidly filtered, rinsed, and inoculated to 500 mL Erlenmeyer flasks filled with 35 mL of acetate medium. To minimize the stress for the cells, all equipment and solutions were prewarmed to 37°C and the transfer was performed within less than two minutes. Cell growth was determined by measuring the OD_600_ by spectrophotometry at 0, 15, 45, 90, 120 min and then every hour up to 420 min after inoculation. The maximal growth rate was calculated using time-points after 240 min and lag time was calculated as previously described (Enjalbert, Cocaign-Bousquet, Portais, & Letisse, 2015).

### Biofilm

Wild-type, Δ*glgP*, Δ*glgA* cells were grown over night at 37°C in Luria Broth (LB) until the cells entered stationary phase. Cells were transferred to a non-shaking environment at room temperature to induce biofilm formation. OD_600_ of the supernatant was measured every ∼30 min.

### Glycogen content experiments

For the depletion experiment, wild-type cells were grown in M9 media and glucose until OD 0.5-0.8. Cells were rapidly transferred into M9 media without carbon source to initiate starvation. Samples were taken before starvation and 10 min, 30 min and 50 min after starvation. For sampling, 1 mL were taken from the culture and kept on ice. To process the samples, they were centrifuged in a cooled centrifuge at maximum speed for 5 minutes. After centrifugation, 100 μl of BPER were added and the samples were gently shaken for 10 minutes. The samples were again centrifuged for 5 minutes at maximum speed in a cooled centrifuge and the supernatant was transferred to a fresh tube and stored at −20 degrees until further processing. For the assay, 25 μl of the supernatant were hydrolyzed and processed as described in the MAK016 assay kit instructions for colorimetric assays (Sigma-Aldrich). For the replenishment experiment, wild-type cells were grown in M9 media and glucose until OD 0.5-0.8. After a starvation period of 30 min in M9 without carbon source, fructose (200g/L) and thiamine-HCl were added (alternative carbon source to avoid convolution with the assay). The samples were taken before the addition of fructose and 2 min, 5 min and 30 min after the addition and processed as described above. The glycogen content was measured with a fluorometric method as described in the MAK016 assay kit instructions (Sigma-Aldrich).

### Microfluidics setup

The custom method of delivering controlled fluctuating nutrient environments is described in previous work (Nguyen, Fernandez, et al., 2019). In brief, microfluidic channels with a depth of 60 μm were cast in polydimethylsiloxane (PDMS). Each PDMS (Sylgard 184; Dow Corning) device was bonded to a glass slide by plasma treating each interacting surface for at least 1 min, and the assembled chip then incubated for at least 2 h at 80°C. The morning of each experiment, bonded channels were cooled to room temperature and then treated with a 1:10 dilution of poly-L-lysine (Sigma catalog no. P8920) in Milli-Q water. This treatment enhanced cell attachment but did not affect growth rate. Wild-type YFP cells and mutant CFP cells were grown over night in M9 medium with glucose and ampicillin. The cultures were then inoculated in fresh M9 medium with glucose and ampicillin. After growing until OD 0.5 – 1.0, the cells were filtered and transferred to a 1:8 diluted M9 medium without glucose (starvation medium) to a final OD of 0.2. Afterwards, the cells were inoculated into the microchannel. Connecting all inputs and outputs to the microchannel took about 10–15 min, allowing ample time for cells to settle and attach to the glass surface within each microchannel before flow was established. By the onset of the fluctuating nutrient signal, cells were without carbon for at least 30 min. The fluctuating signal delivered 5 min periods of carbon free MOPS medium (Teknova) separated by 30 s periods of 2% LB medium (100% LB diluted in MOPS medium). The same 2% LB medium was steadily delivered to the non-fluctuating control environment.

### Image acquisition and analysis

Bacterial growth within the microfluidic channels was imaged using phase contract microscopy with a Nikon Eclipse Ti microscope, equipped with an Andor Zyla sCMOS camera (6.5 μm per pixel) at 60x magnification (40x objective with 1.5x amplification), for a final image resolution of 0.1083 μm per pixel. Each position was repeatedly imaged every 3 min. Image series were processed using a custom MATLAB particle tracking pipeline, which identified individual particles based on pixel intensity and measured particle parameters, such as width and length. These size parameters were used to (1) filter particles that were associated with multiple cells or cells in close proximity to another and (2) approximate the volume of each single cell as a cylinder with hemispherical caps. The approximated volumes were then used to compute instantaneous single-cell growth rates in terms of volume doublings per hour. Using *V(t+*Δ*t)* = *V(t)* · 2^μΔ*t*^, we calculated μ between each pair of time points, associating the resulting μ with the latter of the two time points.

## Data and Code Availability

All data and code used for figure generation are available in **Supplementary Data** or at https://github.com/karsekar/glycogen-starvation.

## Supplementary Information

**Table S1.**
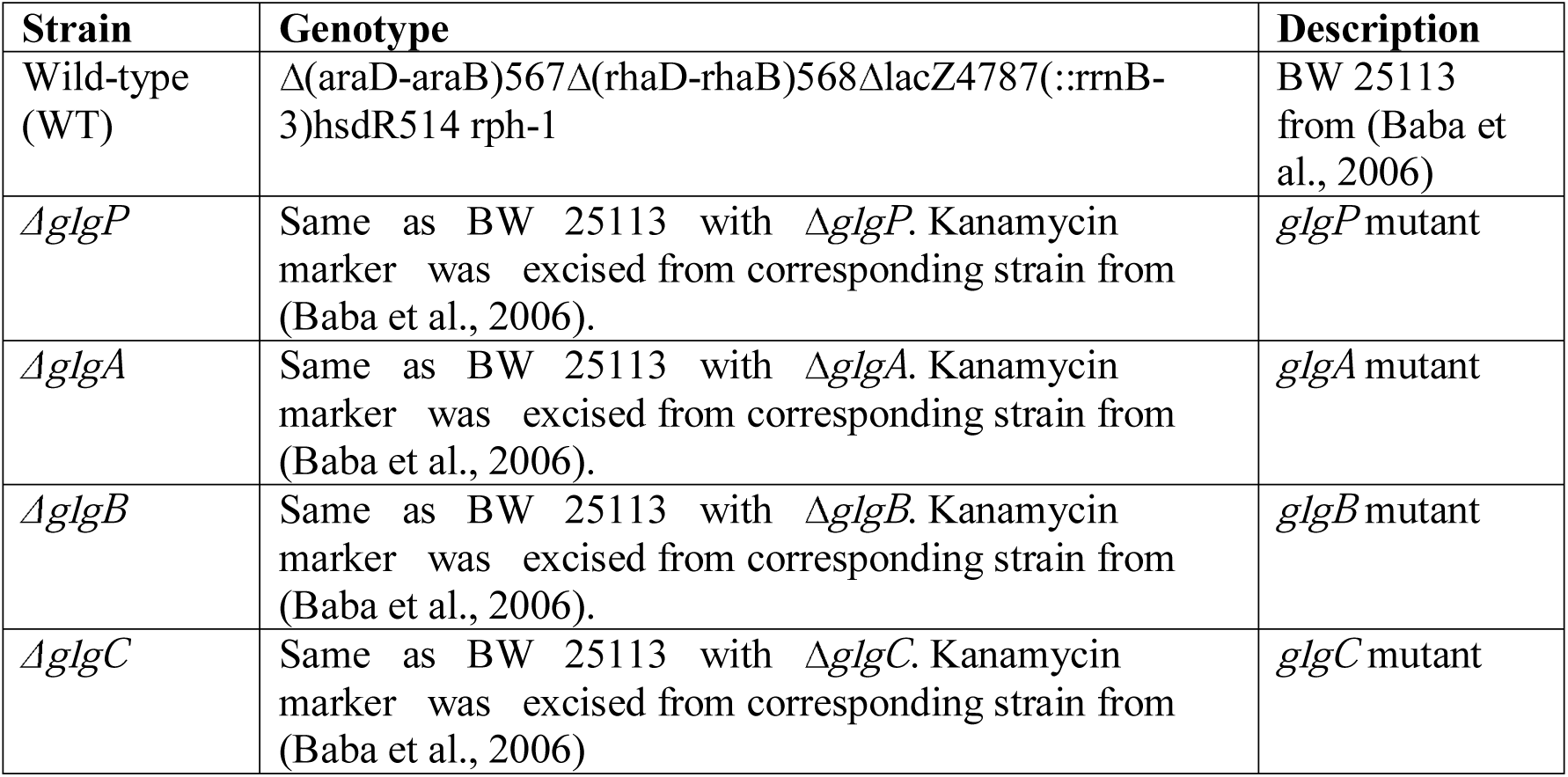
All strains used in this study.

**Table S2.** All plasmids used in this study.

| Plasmid | Description | Reference |
| --- | --- | --- |
| YFP | Plasmid used to provide constitutive expression of yellow fluorescence protein (pRSET-B YFP AddgeneID: #108856). | (Sarabipour, King, & Hristova, 2014) |
| CFP | Plasmid used to provide constitutive expression of cyan fluorescein protein (pRSET-B CFP AddgeneID: #108858). | (Sarabipour et al., 2014) |

**Figure S1.**
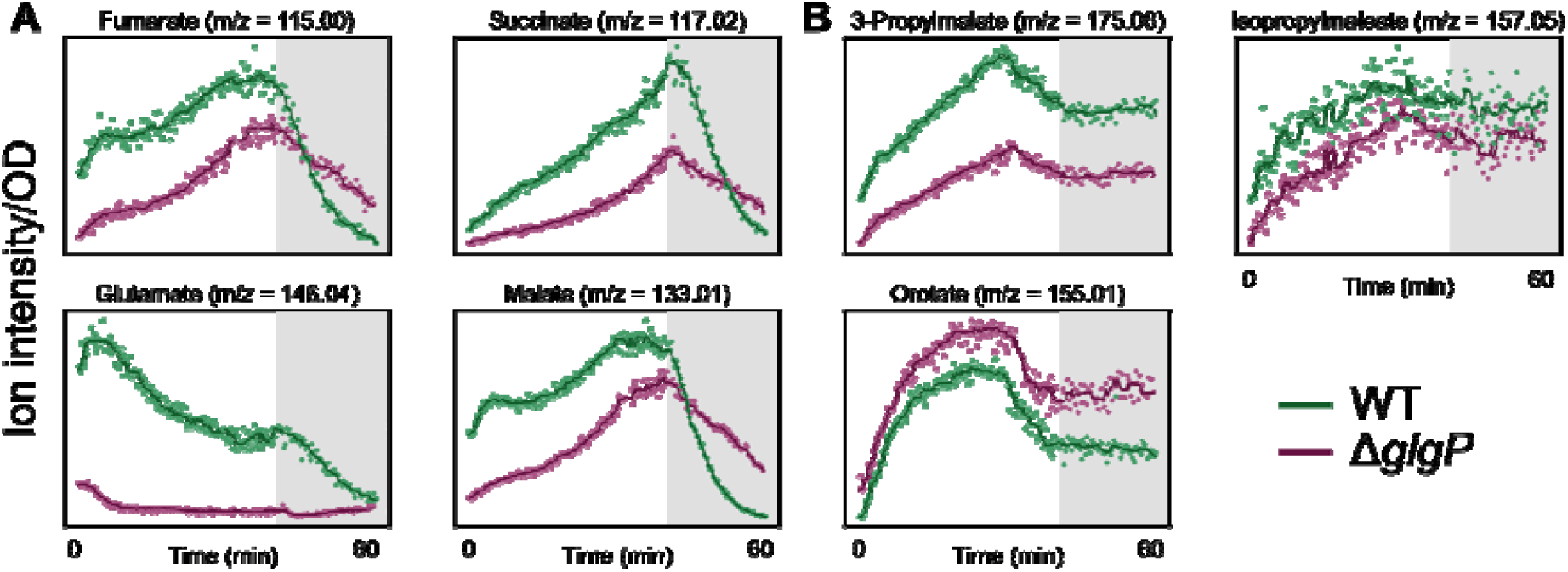
Additional metabolic traces during glucose depletion. (**A**) Various metabolites in the TCA cycle deplete after glucose is depleted out of the media regardless of glycogen capability. Two strains, WT (Wild-type, green) and a *glgP* mutant (purple) are shown, as indicated by the color. Dots indicate ion intensity measurement normalized to initial OD. Gray area indicates the time period after glucose depletion. Solid lines are a moving average filter of the measured ion intensity. (**B**) Traces of exemplary ions that remain constant after glucose depletion for both wild-type and *glgP*.

**Figure S2.**
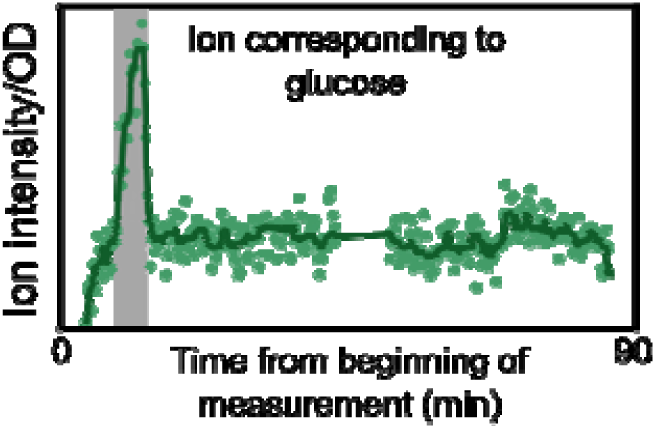
The ion corresponding to glucose rises and falls with the availability of glucose feed. *E. coli* was grown to OD 0.8-1.2, starved for 30 min without glucose and fed a constant glucose supply. The glucose was supplied with a pump at a rate of 8 mmol/g/h for two strains, WT (wild-type, green) and a *glgP* mutant (purple). After 5 min of glucose application, the glucose feed was ceased. Throughout the feed, real-time metabolomics measurement was performed, and data is shown for ions corresponding to glucose as in Figure 3. Dots indicate ion intensity measurement normalized to initial OD. Gray area indicates the time period where glucose was supplied. Solid lines are a moving average filter of the measured ion intensity.

**Figure S3.**
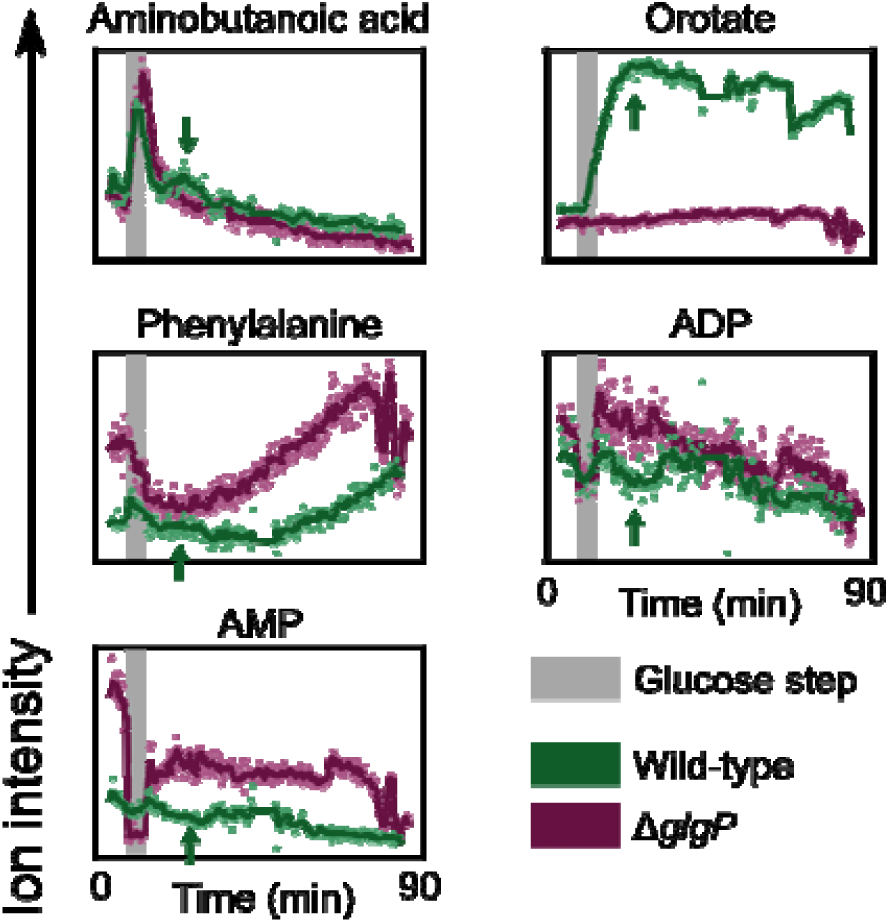
Additional ion traces for glucose step experiment. *E. coli* was grown to OD 0.8-1.2, starved for 30 min without glucose and fed a constant glucose supply. The glucose was supplied with a pump at a rate of 8 mmol/g/h for two strains, WT (wild-type, green) and a *glgP* mutant (purple). After 5 min of glucose application, the glucose feed was ceased. Throughout the feed, real-time metabolomics measurement was performed, and data is shown for ions corresponding to metabolites (aminobutanoic acid, orotate, phenylalanine) and energy charges (AMP and ADP) as in Figure 3. Dots indicate ion intensity measurement normalized to initial OD. Gray area indicates the time period where glucose was supplied. Solid lines are a moving average filter of the measured ion intensity.

## Author contributions

K.S. conceived the project. All authors designed the experiments. K.S., S.M.L., J.N. and A.G. developed the methodology, executed the experiments, and analyzed the data. U.S. and R.S. supervised the work. K.S., S.M.L., and J.N. wrote the manuscript. All authors reviewed and approved the manuscript.

## Acknowledgements

We thank the Sauer laboratory members for useful comments and feedback on the manuscript. We additionally thank T. Conway for discussions, and C. Gao for important advice on the image analysis.

